# Contributions of distinct auditory cortical inhibitory neuron types to the detection of sounds in background noise

**DOI:** 10.1101/2021.06.12.448208

**Authors:** Anna A. Lakunina, Nadav Menashe, Santiago Jaramillo

**Author notes:** **Corresponding author:** Santiago Jaramillo. 1254 University of Oregon. Eugene, OR 97403. **Author contributions:** AAL and SJ conceived the project and designed the experiments. AAL and NM conducted the experiments. AAL and SJ analyzed the data. AAL and SJ wrote the paper.

## Abstract

The ability to separate background noise from relevant acoustic signals is essential for appropriate sound-driven behavior in natural environments. Examples of this separation are apparent in the auditory system, where neural responses to behaviorally relevant stimuli become increasingly noise-invariant along the ascending auditory pathway. However, the mechanisms that underlie this reduction in responses to background noise are not well understood. To address this gap in knowledge, we first evaluated the effects of auditory cortical inactivation on mice of both sexes trained to perform a simple auditory signal-in-noise detection task, and found that outputs from the auditory cortex are important for the detection of auditory stimuli in noisy environments. Next, we evaluated the contributions of the two most common cortical inhibitory cell types, parvalbumin-expressing (PV^+^) and somatostatin-expressing (SOM^+^) interneurons, to the perception of masked auditory stimuli. We found that inactivation of either PV^+^ or SOM^+^ cells resulted in a reduction in the ability of mice to determine the presence of auditory stimuli masked by noise. These results indicate that a disruption of auditory cortical network dynamics by either of these two types of inhibitory cells is sufficient to impair the ability to separate acoustic signals from noise.

**Significance Statement:** Appropriate behavior in a natural environment relies on the ability to separate background noise from relevant signals. We found that auditory cortical inhibitory neurons play a causal role in separating environmental noise from behaviorally relevant auditory signals. These results advance our understanding of the computations performed by the auditory system to decompose and analyze acoustic stimuli in the presence of noise.

## Introduction

The ability to separate background noise from relevant signals is essential for appropriate behavior in natural environments. There is clear evidence that such a separation occurs in the auditory system, where the neural representation of behaviorally relevant stimuli (*e*.*g*., intraspecies vocalizations) becomes increasingly noise-invariant along the ascending auditory pathway (Rabinowitz et al., 2013; Schneider and Woolley, 2013; Carruthers et al., 2015). A potential strategy used by the auditory system to separate signals from noise is to emphasize acoustic features that are common in behaviorally relevant stimuli while suppressing features characteristic of background noise. For instance, the auditory system can make use of the statistics of environmental noise, which tends to be broadband and comodulated, to filter it out (Nelken et al., 1999). In this study, we examined the contribution of cortical inhibition to the implementation of such computation. The sound-evoked response of a large subset of auditory cortical neurons is suppressed as the bandwidth of the stimulus increases, with cortical inhibition being proposed as the possible mechanism by which this suppression takes place (Rauschecker and Tian, 2004; O’Connor et al., 2010; Li et al., 2019). In particular, one class of cortical inhibitory interneurons, those that express somatostatin (SOM^+^), appears to play a unique role in mediating lateral inhibition (Kato et al., 2017) and suppressing neural responses to noise-like broadband stimuli (Lakunina et al., 2020), suggesting a unique role for these cells in filtering background noise. As distinct cortical inhibitory interneuron types are known to have different functions in sensory cortices (Adesnik et al., 2012; Natan et al., 2015; Phillips et al., 2017), we sought to determine which sources of cortical inhibition are important for acoustic signal detection in noise. Because of the unique contributions of SOM^+^ cells to suppressing neural responses to broadband noise, we hypothesized that these cells play a major role in the detection of masked stimuli, while other inhibitory interneuron types, such as parvalbumin-expressing (PV^+^) cells, play only a minimal role in this process. Our results, however, demonstrate that perturbation of either PV^+^ or SOM^+^ cells yields consistent deficits in signal detection in noise, suggesting that a disruption of the cortical network dynamics by either cell type is sufficient to impair the ability of the auditory cortex to separate acoustic signals from noise.

## Materials and Methods

### Animal subjects

Data from a total of 49 transgenic adult mice of both sexes, age 3-9 months, were used in this study. Optical fibers were implanted in 31 mice for optogenetic manipulation during behavior (8 PV::ChR2, 13 PV::ArchT, 10 SOM::ArchT), and a headbar was implanted in one PV::ArchT mouse for electrophysiological recordings. PV-Cre and SOM-Cre driver lines from The Jackson Laboratory (JAX #008069 and #013044) were crossed with LSL-ChR2 mice (JAX #012569) or LSL-ArchT mice (JAX #021188) to produce mice expressing channelrhodopsin-2 (ChR2) or archaerhodopsin (ArchT, (Han et al., 2011)) in parvalbumin-expressing (PV^+^) or somatostatin-expressing (SOM^+^) inhibitory interneurons. All mice were housed in groups of same-sex littermates, and continued to be housed together after surgery whenever possible. Mice were housed in a 12:12 hour light-dark cycle, and all experiments were carried out during the dark period, when mice are most active. All procedures were carried out in accordance with National Institutes of Health Standards and were approved by the University of Oregon Institutional Animal Care and Use Committee.

### Behavioral task

The signal detection task was carried out inside single-walled sound-isolation boxes (IAC Acoustics, Naperville, IL). Mice were water-restricted to motivate them to perform the task. Mice were weighed and their health checked after each behavioral session, and they were provided with a water supplement if their weight was below 80% of their baseline. Except for these supplements, access to water was restricted to the time of the task during experimental days. Free water was provided on days with no experimental sessions. Behavioral data was collected using the *taskontrol* software platform (www.github.com/sjara/taskontrol) written in the Python programming language (www.python.org). Mice initiated each trial by poking their noses into the center port of a three-port behavior chamber. After a silent delay of random duration (200–400 ms, uniformly distributed), a sound was presented until the animal withdrew from the port, to a maximum of 500 ms. Animals were required to then choose one of the two side ports to obtain a reward (2 *µ*l of water). Twenty mice were trained to report the presence of a tone by going right and the absence of a tone by going left, while eleven mice were trained to report the presence of a tone by going left and the absence of a tone by going right. At the end of the sound stimulus, the mice had 4 seconds to make their choice and go to one of the reward ports. If a mouse did not respond in this period of time, the trial was aborted and not considered during data analysis. Aborted trials made up less than 2% of trials in every session (the median across animals was below 0.5% of trials), so their exclusion is unlikely to affect the interpretation of the behavioral data.

Sounds consisted of a noise stimulus acting as a masker and an 8 kHz pure tone stimulus acting as the signal to be detected. Two distinct bandwidths (0.25 octaves and white noise) were used for the noise maskers. The 0.25 octave masker was produced by bandpass filtering white noise. The masker was sinusoidally amplitude-modulated at a rate of 8 Hz and depth of 100%, while the pure tone was unmodulated. The pure tone stimulus was present in 50% of trials. When the pure tone was present, it appeared simultaneously with the noise, turning on and off at the same time. The signal-to-noise ratio (SNR) was varied by changing the amplitude of the signal. Within a behavioral session, we used 3 distinct SNRs (10, 15, and 20 dB) for the pure tone signal. The intensity of all frequency components of the noise was a single value that varied on a per-trial basis from 30 to 40 dB-SPL during the initial training, but was fixed for all trials at 40 dB-SPL during testing. Each behavioral session lasted 60-90 min. Mice took an average of 19 days (range: 10 to 42 days) from the beginning of training to reach a performance of 70% correct in the simplest version of the signal detection task, where loud pure tone signals were masked by white noise. The data presented in Fig. 1 contains at least 5 sessions of pre-implantation performance from each mouse.

**Figure 1:**
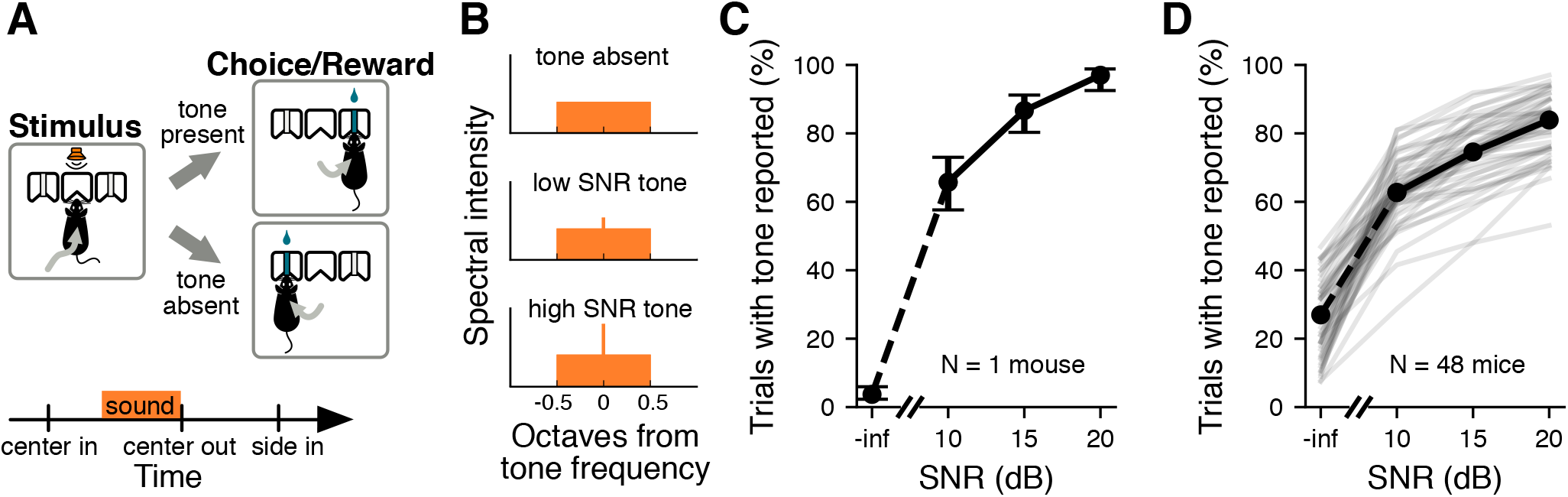
Performance in a signal detection task. **A**, Schematic of the signal detection task. Mice had to correctly report the presence or absence of a pure tone signal to obtain a reward. **B**, Example frequency spectra of sounds used during the signal detection task. **C**, Example psychometric curve showing performance of one mouse during one behavior session. Performance is averaged over all masker bandwidths presented. Error bars show 95% confidence intervals. **D**, Median psychometric curve (black line) for mice trained in the signal detection task (*N* = 48 mice). Psychometric curves for individual mice are shown in gray. Trials are pooled across multiple sessions. Performance is averaged over all masker bandwidths presented.

### Optogenetic stimulation in awake mice

Optical fibers (CFML12U-20, 200 *µ*m core diameter, ThorLabs) were cleaved and etched with hydrofluoric acid for 40 min to obtain a cone-shaped tip. Each optical fiber was glued to a metal guide tube that helped secure the fiber ferrule to the skull. Before implantation, optical fibers were connected to a laser (445 nm for activating ChR2 or 520 nm for activating ArchT) built in-house and the light output calibrated using a PM100D power meter (ThorLabs). To implant the fibers, animals were anesthetized with isoflurane through a nose-cone on a stereotaxic apparatus before the optical fibers were implanted bilaterally in the auditory cortex at 2.8 mm posterior to bregma, 4.4 mm from midline, and 0.5 mm from brain surface. This location corresponds to the dorsal region of auditory cortex, right above primary auditory cortex (Fig. 2). Laser illumination therefore was concentrated mostly on the primary auditory cortex (AUDp) with light likely extending to parts of the dorsal (AUDd) and ventral (AUDv) regions given the laser powers used (see below). We confirmed that the fiber locations targeted the auditory cortex by imaging brain slices postmortem.

**Figure 2:**
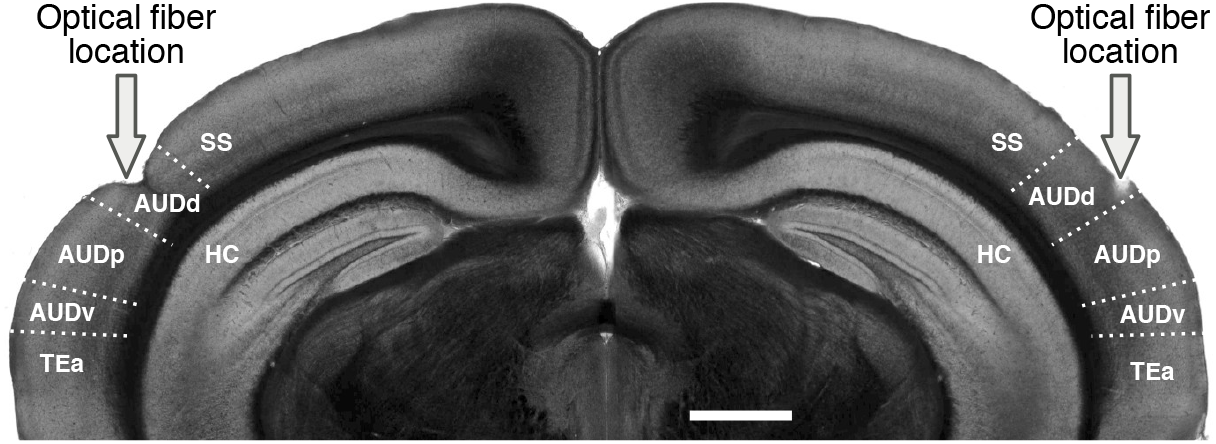
Location of optical fibers. Coronal brain slice showing the location of optical fibers implanted above the auditory cortex, such that light was concentrated on the primary field of the auditory cortex (AUDp). The primary field is surrounded by a dorsal and a ventral fields (AUDd and AUDp). Laser powers were chosen such that areas outside the auditory cortex were unlikely to be affected (SS: somatosensory cortex, TEa: temporal association area, HC: hippocampus). Scale = 1 mm.

During a behavioral session, the laser was connected to the implanted optical fibers with flexible fiber optic patch cables (MFP 200/240/900-0.22 2m FC-MF1.25, Doric Lenses). The cables did not impede the mouse’s movement, and 2-4 sessions were run prior to the experimental sessions to acclimate the mouse to the fibers and ensure it could still perform the task. The laser power used was 3 mW at the tip for PV::ChR2 animals and 10 mW at the tip for PV::ArchT and SOM::ArchT animals. These powers have previously been shown to restrict illumination to the auditory cortex, while limiting effects on firing rate in other cortical areas (Weible et al., 2014a,b). For instance, in PV::ChR2 mice and a blue laser power of 200 mW/mm^2^ (about twice of that used in our ChR2 experiments), significant suppression of activity extended to about 1500 *µ*m from the fiber tip (Weible et al., 2014a). Similarly, in animals expressing Arch, a green laser power of 300 mW/mm^2^ (about the same used in our experiments) resulted in suppression of activity at about the same distance as in the ChR2 experiments (Weible et al., 2014b). Laser was presented in 25% of trials. The laser turned on at sound onset and turned off 100 ms after the sound ended or when the mouse entered one of the reward ports, whichever came first. For each mouse, 8 sessions were performed with laser stimulation directed at the auditory cortex. In between these experimental sessions, we interleaved four additional control sessions, denoted as “laser directed away”, with the patch cables attached to the implant, pointing to the edge of the head cover, but not connected to the optical fibers. In these additional control sessions, animals could see the laser light to the same extent as during laser stimulation sessions, but the laser did not reach the brain. Mice were excluded from analysis if they had fewer than three sessions with a performance of at least 60% correct (1 PV::ChR2 and 2 SOM::ArchT mice excluded). For the included mice, only sessions where the mouse had a performance of at least 60% correct trials were included in the data analysis. The median number of excluded session was 0.5 for PV::ChR2, 0 for PV::ChR2, and 3.5 for SOM::ArchT mice. To quantify the effects of the laser on the timing of the animals’ motor output, we measured the time mice spent sampling the sound, defined as the amount of time between sound onset and withdrawal from the center port, as well as the time mice spent travelling to the reward port, defined as the amount of time between withdrawal from the center port and entry in a reward port.

### Neural recordings

To verify that our laser illumination effectively perturbed inhibitory auditory cortical interneurons, a PV::ArchT mouse was surgically implanted with a head-bar to allow for head-fixed extracellular recordings. Bilateral craniotomies (centered at 2.8 mm posterior to bregma and 4.4 mm from midline) were performed to allow for acute recordings from the auditory cortex. The animal was monitored and recovered fully before electrophysiological experiments.

Electrical signals were collected using an RHD2000 acquisition system (Intan Technologies, Los Angeles, CA) and OpenEphys software (www.open-ephys.org), using silicon probe electrodes (A4×2-tet configuration from NeuroNexus, Ann Arbor, MI). During the experiment, the awake mouse was head-fixed and the probe was lowered vertically into the auditory cortex until spikes were detected. We recorded at multiple depths on each penetration, with recording sites typically 100-150 *µ*m apart to avoid recording from the same cells twice.

Cortical PV^+^ cells were inactivated during the presentation of 1000 ms sounds with 1300 ms light pulses (light onset was 100 ms before sound onset, light offset was 200 ms after sound offset). Green light (520 nm wavelength) was delivered via an optical fiber (200 *µ*m diameter) attached to the silicon probe electrodes with the tip about 900 *µ*m from the topmost recording sites. Light power was 5 mW at the fiber tip and was presented for 50% of trials, with laser and non-laser trials randomly interleaved.

Spiking activity was detected by applying a threshold (40-45 *µ*V) to the bandpass-filtered signals (300 to 6000 Hz). Spiking activity of single units was isolated offline using the automated expectation maximization clustering algorithm Klustakwik (Kadir et al., 2014). Isolated clusters were only included in the analysis if less than 2% of inter-spike intervals were shorter than 2 ms. We also calculated a spike quality index, defined as the ratio between the peak amplitude of the waveform and the average variance, calculated using the channel with the largest amplitude. Cells were only included in the analysis if they had a spike quality index greater than 2.5.

### Histology

At the conclusion of each experiment, animals were deeply anesthetized with euthasol and perfused through the heart with 4% paraformaldehyde. Brains were extracted and left in 4% paraformaldehyde for at least 24 hours before slicing. Brains were sliced under phosphate-buffered saline on a vibratome with a slice thickness of 100 *µ*m. Brain slices were imaged with a fluorescent microscope (Axio Imager 2, Carl Zeiss) with a 2.5x objective. Expression of transgenes was verified by the presence of fluorescence in the auditory cortex and by the general pattern of fluorescence expected throughout the brain for each mouse line. To determine the locations of our fiber implants, we manually registered each brain slice to the corresponding coronal section in the Allen Mouse Common Coordinate Framework (Common Coordinate Framework v.3, c 2015 Allen Institute for Brain Science, Allen Brain Atlas API, available from http://brain-map.org/api/index.html).

### Analysis of behavioral data

To quantify how well a subject was able to detect the presence of a pure tone signal, we calculated the subject’s sensitivity index (*d*′) from the hit rate and false alarm rate, as follows:

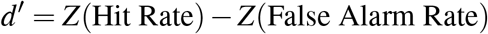

*Z* in this case is the inverse cumulative distribution function of the standard normal. Hit rates were calculated as the percentage of trials containing the signal where the mouse correctly reported the signal’s presence. False alarm rates were calculated as the percentage of trials not containing the signal where the mouse incorrectly reported the signal’s presence.

### Statistical Analysis

To test for the effects of manipulating a particular cell type, we used the non-parametric Wilcoxon signed-rank test because the data were not expected to be normally distributed or of equal variance. To test for differences in the effect of inactivating PV^+^ or SOM^+^ cells, we used the non-parametric Wilcoxon rank-sum test. To test for correlations between two continuous variables, we computed a correlation coefficient using least-squares linear regression. Further details on the comparisons made and statistical tests used can be found in Table 1.

**Table 1:** Summary of statistical analyses.

| Figure | Measurement | Comparison | Statistical test | Test results |
| --- | --- | --- | --- | --- |
| 3D | $d'$ | Without laser presentation vs. with laser presentation ($N = 8$ PV::ChR2 mice) | Wilcoxon signed-rank test | $p = 0.0117$<br>$T = 0$ |
| 3E | Hit rate (%) | Without laser presentation vs. with laser presentation ($N = 8$ PV::ChR2 mice) | Wilcoxon signed-rank test | $p = 0.0117$<br>$T = 0$ |
| 3F | False alarm rate (%) | Without laser presentation vs. with laser presentation ($N = 8$ PV::ChR2 mice) | Wilcoxon signed-rank test | $p = 0.0251$<br>$T = 2$ |
| 3G | Laser-induced change in $d'$ | Laser directed at AC vs. laser directed away from AC ($N = 8$ PV::ChR2 mice) | Correlation coefficient | $p = 0.593$<br>$r = 0.224$ |
| 3H | Laser-induced change in hit rate (%) | Laser directed at AC vs. laser directed away from AC ($N = 8$ PV::ChR2 mice) | Correlation coefficient | $p = 0.710$<br>$r = 0.157$ |
| 3I | Laser-induced change in false alarm rate (%) | Laser directed at AC vs. laser directed away from AC ($N = 8$ PV::ChR2 mice) | Correlation coefficient | $p = 0.589$<br>$r = 0.227$ |
| 5A | Sampling time (ms) | Without laser presentation vs. with laser presentation ($N = 8$ PV::ChR2 mice) | Wilcoxon signed-rank test | $p = 0.0925$<br>$T = 6$ |
| 5B | Time to reward (ms) | Without laser presentation vs. with laser presentation ($N = 8$ PV::ChR2 mice) | Wilcoxon signed-rank test | $p = 0.779$<br>$T = 16$ |
| 5C | Laser-induced change in sampling time (ms) | Laser directed at AC vs. laser directed away from AC ($N = 8$ PV::ChR2 mice) | Correlation coefficient | $p = 0.675$<br>$r = 0.177$ |
| 5D | Laser-induced change in time to reward (ms) | Laser directed at AC vs. laser directed away from AC ($N = 8$ PV::ChR2 mice) | Correlation coefficient | $p = 0.686$<br>$r = -0.171$ |
| 6C | $d'$ | Without laser presentation vs. with laser presentation ($N = 13$ PV::ArchT mice) | Wilcoxon signed-rank test | $p = 0.0046$<br>$T = 5$ |
| 6D | Hit rate (%) | Without laser presentation vs. with laser presentation ($N = 13$ PV::ArchT mice) | Wilcoxon signed-rank test | $p = 0.0330$<br>$T = 15$ |
| 6E | False alarm rate (%) | Without laser presentation vs. with laser presentation ($N = 13$ PV::ArchT mice) | Wilcoxon signed-rank test | $p = 0.861$<br>$T = 43$ |
| 6F | Laser-induced change in $d'$ | Laser directed at AC vs. laser directed away from AC ($N = 13$ PV::ArchT mice) | Correlation coefficient | $p = 0.609$<br>$r = 0.157$ |
| 6G | Laser-induced change in hit rate (%) | Laser directed at AC vs. laser directed away from AC ($N = 13$ PV::ArchT mice) | Correlation coefficient | $p = 0.657$<br>$r = -0.136$ |
| 6H | Laser-induced change in false alarm rate (%) | Laser directed at AC vs. laser directed away from AC ($N = 13$ PV::ArchT mice) | Correlation coefficient | $p = 0.981$<br>$r = -0.0073$ |
| 6K | $d'$ | Without laser presentation vs. with laser presentation ($N = 10$ SOM::ArchT mice) | Wilcoxon signed-rank test | $p = 0.0051$<br>$T = 0$ |
| 6L | Hit rate (%) | Without laser presentation vs. with laser presentation ($N = 10$ SOM::ArchT mice) | Wilcoxon signed-rank test | $p = 0.0051$<br>$T = 0$ |
| 6M | False alarm rate (%) | Without laser presentation vs. with laser presentation ($N = 10$ SOM::ArchT mice) | Wilcoxon signed-rank test | $p = 0.878$<br>$T = 26$ |
| 6N | Laser-induced change in $d'$ | Laser directed at AC vs. laser directed away from AC ($N = 10$ SOM::ArchT mice) | Correlation coefficient | $p = 0.546$<br>$r = 0.218$ |
| 6O | Laser-induced change in hit rate (%) | Laser directed at AC vs. laser directed away from AC ($N = 10$ SOM::ArchT mice) | Correlation coefficient | $p = 0.0494$<br>$r = 0.633$ |
| 6P | Laser-induced change in false alarm rate (%) | Laser directed at AC vs. laser directed away from AC ($N = 10$ SOM::ArchT mice) | Correlation coefficient | $p = 0.0725$<br>$r = 0.590$ |
| 6 | Laser-induced change in $d'$ | Laser directed at AC in PV::ArchT mice ($N=13$) vs. SOM::ArchT mice ($N=10$) | Wilcoxon rank-sum test | $p = 0.321$<br>$T = 0.992$ |
| 8A | Sampling time (ms) | Without laser presentation vs. with laser presentation ($N = 13$ PV::ArchT mice) | Wilcoxon signed-rank test | $p = 0.0652$<br>$T = 15.5$ |
| 8B | Laser-induced change in sampling time (ms) | Laser directed at AC vs. laser directed away from AC ($N = 13$ PV::ArchT mice) | Correlation coefficient | $p = 0.0017$<br>$r = 0.780$ |
| 8C | Time to reward (ms) | Without laser presentation vs. with laser presentation ($N = 13$ PV::ArchT mice) | Wilcoxon signed-rank test | $p = 0.0029$<br>$T = 1$ |
| 8D | Laser-induced change in time to reward (ms) | Laser directed at AC vs. laser directed away from AC ($N = 13$ PV::ArchT mice) | Correlation coefficient | $p = 0.901$<br>$r = 0.0383$ |
| 8E | Sampling time (ms) | Without laser presentation vs. with laser presentation ($N = 10$ SOM::ArchT mice) | Wilcoxon signed-rank test | $p = 0.0050$<br>$T = 0$ |
| 8F | Laser-induced change in sampling time (ms) | Laser directed at AC vs. laser directed away from AC ($N = 10$ SOM::ArchT mice) | Correlation coefficient | $p = 0.901$<br>$r = 0.0453$ |
| 8G | Time to reward (ms) | Without laser presentation vs. with laser presentation ($N = 10$ SOM::ArchT mice) | Wilcoxon signed-rank test | $p = 0.203$<br>$T = 15$ |
| 8H | Laser-induced change in time to reward (ms) | Laser directed at AC vs. laser directed away from AC ($N = 10$ SOM::ArchT mice) | Correlation coefficient | $p = 0.470$<br>$r = 0.259$ |

## Results

### Detection of acoustic signals in noise

To investigate the neuronal mechanisms involved in the detection of acoustic signals in background noise, we trained mice to perform a signal detection task where they had to report the presence or absence of a pure tone signal (8 kHz) immersed in an amplitude modulated noise masker (Fig. 1A, B). Mice successfully learned this task, and their ability to detect the pure tone signal was highest for trials where the signal-to-noise ratio (SNR) was the largest (Fig. 1C,D).

Consistent with the effect of comodulation masking release (Schooneveldt and Brian Moore, 1989; Sollini and Chadderton, 2016), sensitivity to the pure tone signal was lower when the noise masker had a narrow bandwidth (0.25 octaves) compared to a broad bandwidth (white noise), despite the greater power contained within the broadband noise masker (median *d*′ = 0.797 across mice for narrowband masker, median *d*′ = 1.190 for broadband, *p* = 4.67 *×* 10^−6^, Wilcoxon signed-rank test, *N* = 48 mice). Hit rates and false alarm rates were both significantly higher when the masker was narrowband rather than broadband (median narrowband hit rate = 81%, median broadband hit rate = 65%, *p* = 3.38 *×* 10^−13^; median narrowband false alarm rate = 46%, median broadband hit rate = 21%, *p* = 5.11 *×* 10^−13^; Wilcoxon signed-rank test). This result indicates that mice reported the presence of tones more often when the masker was narrowband, even when the tone was not present. Yet, because the improvements in hit rate for narrowband masker trials were not as high as the increases in false alarms, the overall performance was lower compared to trials with a broadband masker. Overall, these results demonstrate that mice can successfully perform a signal-in-noise detection task, and that detection performance depends on the conditions of the stimulus (SNR and bandwidth of the masker) as observed in other species (Schneider and Woolley, 2013; von Trapp et al., 2016).

### Inactivation of the auditory cortex impairs detection of signals in noise

To determine the contribution of auditory cortical circuits to the detection of acoustic signals in background noise, we measured the animals’ performance in the task described above while inactivating the auditory cortex bilaterally in a subset of trials using optogenetics. Mice expressing the light-gated ion channel channelrhodopsin-2 (ChR2) in parvalbumin-expressing inhibitory interneurons (PV^+^) were trained in the task. Optogenetic activation of cortical PV^+^ cells using ChR2 has been shown to be a reliable method for reversibly silencing cortical circuits (Sachidhanandam et al., 2013; Glickfeld et al., 2013), allowing us to determine the effects of cortical inactivation on signal detection in noise. To this end, we implanted mice with optical fibers and delivered blue light to the auditory cortex in both hemispheres during the task (Fig. 3A). Histological analysis post-mortem was performed to ensure the fibers were located over the auditory cortex (Fig. 2).

**Figure 3:**
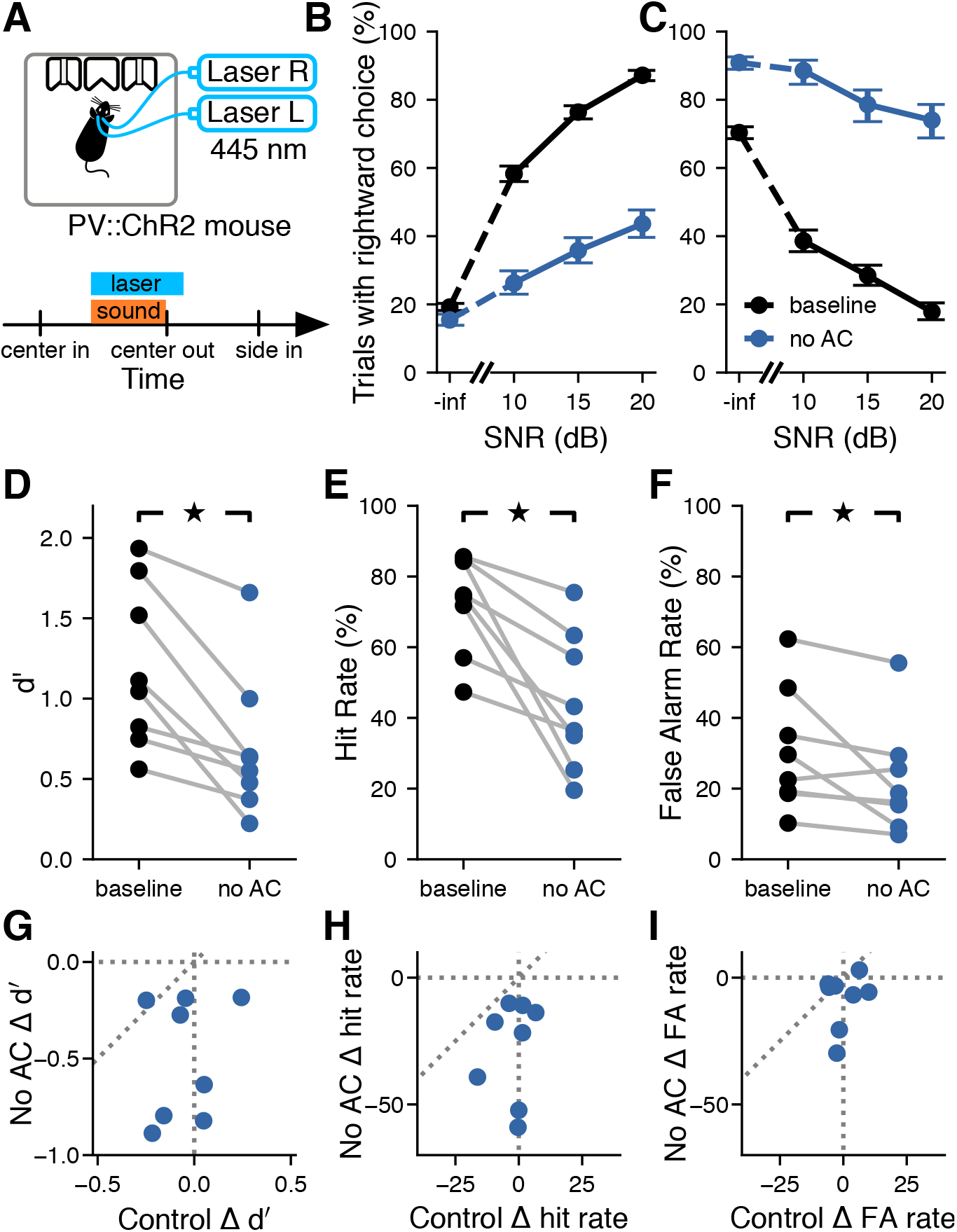
Inactivation of the auditory cortex impaired the detection of signals masked by noise. **A**, Top: optogenetic stimulation in freely-moving PV::ChR2 mice. Laser activation of PV^+^ cells in the auditory cortex silences auditory cortical activity. Bottom: For a random 25% of trials in each session, the laser turned on at the same time as the sound and lasted for 100 ms after the sound turned off. **B**, Example psychometric curve from one PV::ChR2 mouse trained to report the presence of the tone by going to the right (trials pooled across multiple sessions). Activation of auditory cortical PV^+^ cells reduced the probability of the mouse reporting the presence of the tone for all SNRs. Error bars show 95% confidence intervals. **C**, Like B, but for a PV::ChR2 mouse trained to go to the left to report the presence of the tone. As before, activation of auditory cortical PV^+^ cells reduced the probability of the tone being reported, despite the opposite action being required to report tones. **D**, Silencing the auditory cortex during the signal detection task significantly reduced sensitivity to the pure tone signal (*p* = 0.0117, Wilcoxon signed-rank test, *N* = 8 mice). Each pair of points corresponds to one mouse. **E**, Silencing the auditory cortex during the signal detection task significantly reduced the hit rate (*p* = 0.0117, Wilcoxon signed-rank test). **F**, Silencing the auditory cortex during the signal detection task significantly reduced the false alarm rate (*p* = 0.0251, Wilcoxon signed-rank test). **G**, The effect of silencing the auditory cortex (No AC) on *d*′ was not correlated with the effect of visual distraction (Control) on *d*′ (*p* = 0.593, *r* = 0.224, linear correlation coefficient). Each point corresponds to one mouse. **H**, The effect of silencing the auditory cortex on hit rate was not correlated with the effect of visual distraction on hit rate (*p* = 0.157, *r* = 0.710, linear correlation coefficient). **I**, The effect of silencing the auditory cortex on false alarm rate was not correlated with the effect of visual distraction on false alarm rate (*p* = 0.589, *r* = 0.227, linear correlation coefficient).

Bilateral inactivation of the auditory cortex during sound presentation impaired the performance of the mice in the detection task, reducing their sensitivity to the stimulus (Fig. 3D; *p* = 0.0117, Wilcoxon signed-rank test). While the baseline performance of mice differed by bandwidth of the masker, as described in the previous section, the effects of optogenetic manipulation did not depend on masker bandwidth (median change in *d*′ with 0.25 octave masker across mice: −0.751; median change in *d*′ with white noise masker: −0.553; *p* = 0.575, Wilcoxon signed-rank test). Thus, we pooled trials across bandwidths for subsequent analysis. We found that the decrease in performance level during inactivation trials could not be explained simply by visual distraction from the laser used for optogenetics because the effect of the laser on the sensitivity index (*d*′) was significantly larger when the laser was directed at the auditory cortex than when it was directed away (*p* = 0.0173, Wilcoxon signed-rank test). Moreover, we found no correlation between the change in sensitivity index when the laser was directed at the auditory cortex and the change in sensitivity index when the laser was directed away from the auditory cortex (Fig. 3G; *p* = 0.593, *r* = 0.224, linear correlation coefficient).

The decrease in sensitivity index observed during inactivation of the auditory cortex was largely the result of a decrease in hit rates (Fig. 3E; *p* = 0.0117, Wilcoxon signed-rank test). Contrary to the expectation that auditory cortical inactivation would have a smaller effect on the detection of signals with a high SNR, as the task is easier, we found slightly larger effects on the detection of 20 dB SNR signals compared to 10 dB SNR signals (21.5% reduction in hit rate for SNR of 20 dB, 16.9% reduction for 10 dB, *p* = 0.0499, Wilcoxon signed-rank test). Inactivation of the auditory cortex also significantly reduced false alarm rates (Fig. 3F; *p* = 0.0251, Wilcoxon signed-rank test), although this improvement was not sufficient to overcome the decrease in hit rates and improve sensitivity. The combination of a decrease in hit rate and a decrease in false alarm rate indicates that mice reported the presence of the pure tone signal less often when the auditory cortex was inactivated, regardless of whether the signal was present or not. As with the effect on sensitivity, these effects could not be explained by visual distraction, as the changes in hit rate or false alarm rate were not correlated between the two laser location conditions (Fig. 3H,I, hit rate: *p* = 0.157, *r* = 0.710; false alarm rate: *p* = 0.589, *r* = 0.227; linear correlation coefficient).

The observed deficits during auditory cortex inactivation were not related to which port was associated with the presence or absence of the signal, as mice trained to go to opposite sides to report the presence of the signal were all impaired in their ability to detect the signal (Fig. 3B,C). To verify that the estimated deficits in performance during cortical inactivation were not the result of a few sessions with outlier behavior, we quantified the changes in sensitivity index for each behavioral session individually and plotted these next to estimates obtained by pooling trials across sessions (Fig. 4). Even though variability in behavior from session to session is apparent, these observations support the validity of the results obtained from pooling trials across sessions.

**Figure 4:**
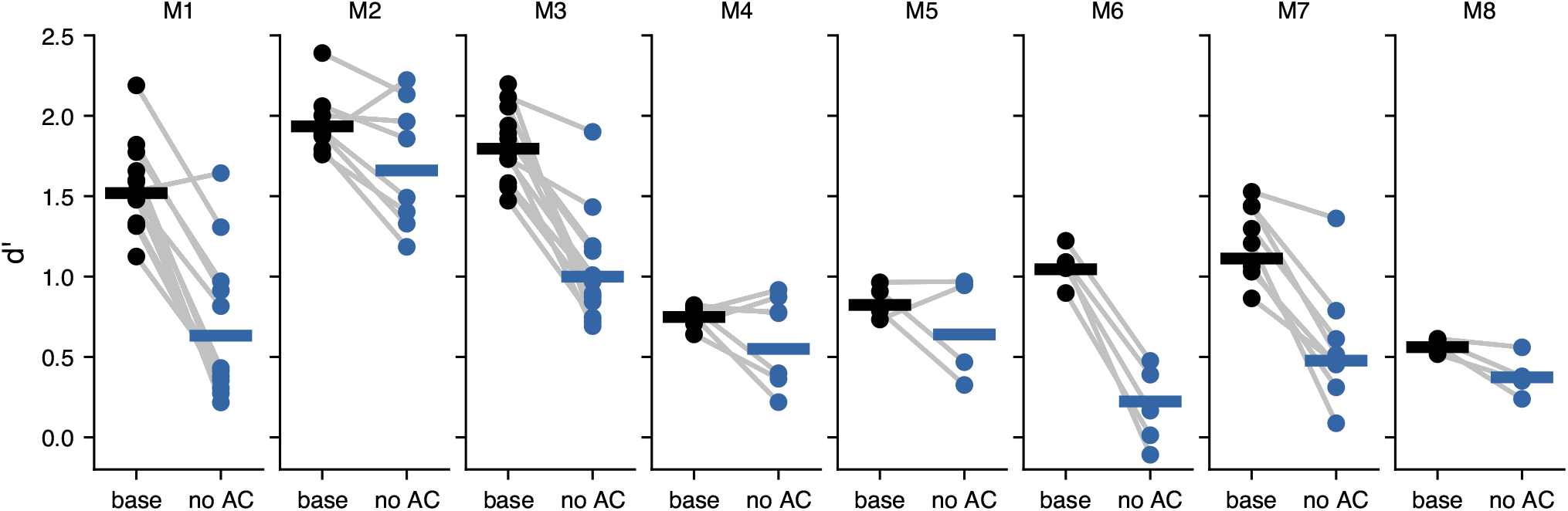
Inactivation of the auditory cortex yielded consistent impairments across sessions. Signal detection performance (*d*′) during each behavioral session (dots) compared to the pooled performance across all trials (horizontal lines) for each PV::ChR2 mouse (M1-M8).

To determine whether inactivation of the auditory cortex resulted in motor deficits during the task, we tested the effects of laser stimulation on the timing of the animals’ motor output. Optogenetic inactivation of the auditory cortex did not significantly affect the amount of time mice spent sampling the sound before making their decision (Fig. 5A; *p* = 0.0925, Wilcoxon signed-rank test), or the amount of time it took mice to reach the reward ports upon leaving the center port (Fig. 5B; *p* = 0.779, Wilcoxon signed-rank test). Changes in sampling time and time to reward were not correlated between conditions where the laser was directed at the auditory cortex or directed away, indicating that the effects of visual distraction were unrelated to any effects brought about by inactivation of the auditory cortex (Fig. 5C,D; sampling time: *p* = 0.675, *r* = 0.177; time to reward: *p* = 0.686, *r* = −0.171; linear correlation coefficient). These results suggests that the effects of auditory cortical inactivation on performance in the signal detection task were the result of a perceptual deficit, not a motor deficit.

**Figure 5:**
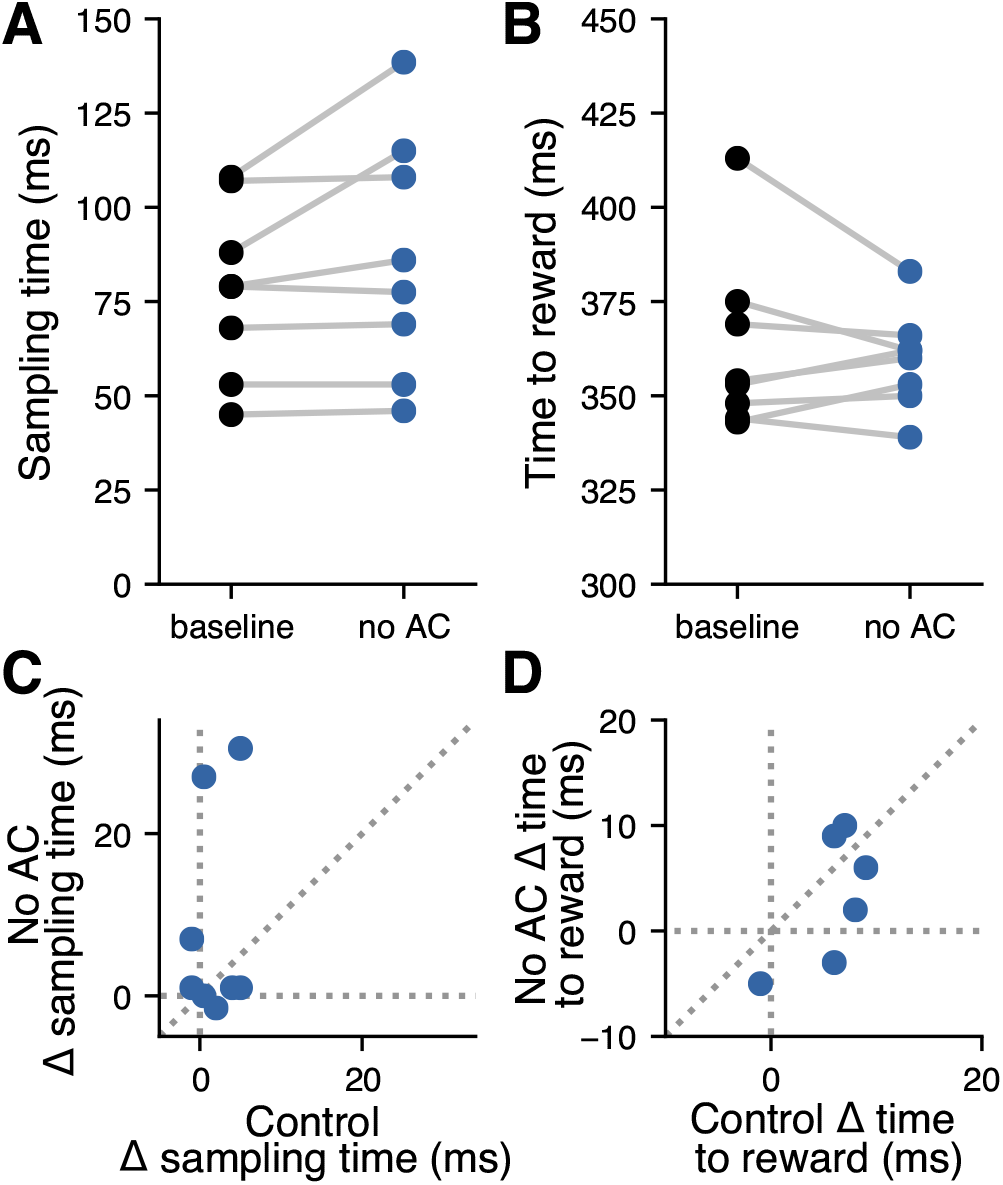
Inactivation of the auditory cortex did not affect timing of behavior. **A**, Silencing the auditory cortex during the signal detection task did not affect the time spent sampling the sound in the center port (*p* = 0.0925, Wilcoxon signed-rank test, *N* = 8 mice). Each pair of points corresponds to one mouse. **B**, Silencing the auditory cortex during the signal detection task did not affect the time spent moving toward the reward port (*p* = 0.779, Wilcoxon signed-rank test). Each point corresponds to one mouse. **C**, The effect of silencing the auditory cortex (No AC) on time spent sampling the sound in the center port was not correlated with the effect of visual distraction (control) on sampling time (*p* = 0.675, *r* = 0.177, linear correlation coefficient). **D**, The effect of silencing the auditory cortex on time spent moving toward the reward port was not correlated with the effect of visual distraction on time spent moving toward the reward port. (*p* = 0.686, *r* = −0.171, linear correlation coefficient).

### Cortical inhibitory neurons contribute to auditory signal detection in noise

To determine the extent to which distinct cortical inhibitory interneuron types are involved in auditory signal detection in noise, we measured the performance of animals in the detection task described above while bilaterally perturbing the activity of either parvalbumin-expressing (PV^+^) or somatostatin-expressing (SOM^+^) neurons from the auditory cortex. We trained mice that express the light-driven outward proton pump archaerhodopsin (ArchT) in either PV^+^ or SOM^+^ cells, allowing us to use light to inactivate these specific neuron classes during behavior. We then implanted these mice with optical fibers to enable delivery of green light to the auditory cortex during sound presentation in a subset of trials (Fig. 6A,I). Histological analysis post-mortem confirmed that the fibers were located over the auditory cortex. To evaluate the physiological effects of our perturbation method, we performed extracellular electrophysiological recordings from the auditory cortex of a naive PV::ArchT mouse while activating ArchT with green light, as in the behavioral experiments. We observed an average increase of 10% in sound-evoked firing rates across the random population of recorded neurons, as expected from a decrease in inhibition in the circuit (*N* = 88 cells, *p* = 1.54 *×* 10^−6^, Wilcoxon signed-rank test). These data suggest that optogenetic inactivation of a particular inhibitory cell type using ArchT perturbs neural activity throughout the auditory cortex.

**Figure 6:**
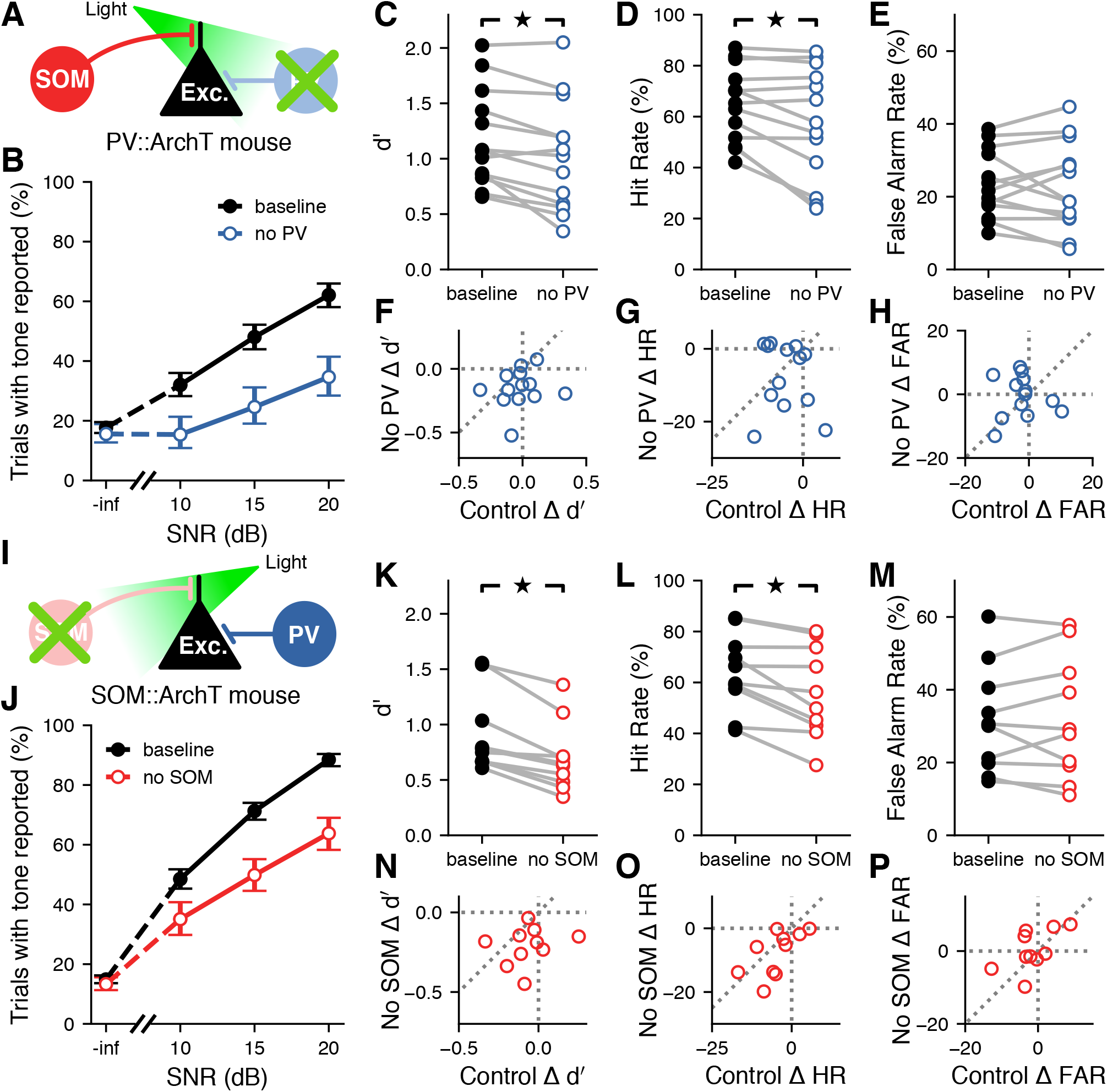
Contributions of distinct inhibitory neuron types to auditory signal detection. **A**, Inactivation of PV^+^ cells during the signal detection task. Laser was presented for a random 25% of trials each session. **B**, Example psychometric curve from one PV::ArchT mouse (trials pooled across multiple sessions). Inactivation of auditory cortical PV^+^ cells decreased the probability of the mouse reporting hearing the signal when it was present, though false alarm rate was unaffected. Error bars show 95% confidence intervals. **C**, Inactivating auditory cortical PV^+^ cells during the signal detection task significantly reduced sensitivity to the signal (*p* = 0.0046, Wilcoxon signed-rank test, *N* = 13 mice). **D**, Inactivating auditory cortical PV^+^ cells during the signal detection task significantly reduced the hit rate (*p* = 0.033, Wilcoxon signed-rank test). **E**, Inactivating auditory cortical PV^+^ cells during the signal detection task did not significantly affect the false alarm rate (*p* = 0.861, Wilcoxon signed-rank test). **F**, The effect of inactivating auditory cortical PV^+^ cells (No PV) on *d*′ was not correlated with the effect of visual distraction (control) on *d*′ (*p* = 0.609, *r* = 0.157, linear correlation coefficient). **G**, The effect of inactivating auditory cortical PV^+^ cells on hit rate was not correlated with the effect of visual distraction on hit rate (*p* = 0.657, *r* = −0.136, linear correlation coefficient). **H**, The effect of inactivating auditory cortical PV^+^ cells on false alarm rate was not correlated with the effect of visual distraction on false alarm rate (*p* = 0.981, *r* = −0.0073, linear correlation coefficient). **I**, Inactivation of SOM^+^ cells during the signal detection task. Laser was presented for a random 25% of trials each session. **J**, Like B, but for one SOM::ArchT mouse. Inactivation of auditory cortical SOM^+^ cells reduced the probability of the mouse reporting hearing the signal when it was present, though false alarm rate was unaffected. **K**, Inactivating auditory cortical SOM^+^ cells during the signal detection task significantly reduced sensitivity to the signal (*p* = 0.0051, Wilcoxon signed-rank test, *N* = 10 mice). **L**, Inactivating auditory cortical SOM^+^ cells during the signal detection task significantly reduced hit rate (*p* = 0.0051, Wilcoxon signed-rank test). **M**, Inactivating auditory cortical SOM^+^ cells during the signal detection task did not significantly affect the false alarm rate (*p* = 0.878, Wilcoxon signed-rank test). **N**, The effect of inactivating auditory cortical SOM^+^ cells (No SOM) on *d*′ was not correlated with the effect of visual distraction (control) on *d*′ (*p* = 0.546, *r* = 0.218, linear correlation coefficient). **O**, The effect of inactivating auditory cortical SOM^+^ cells on hit rate was correlated with the effect of visual distraction on hit rate (*p* = 0.0494, *r* = 0.633, linear correlation coefficient). **P**, The effect of inactivating auditory cortical SOM^+^ cells on false alarm rate was not correlated with the effect of visual distraction on false alarm rate (*p* = 0.0725, *r* = 0.590, linear correlation coefficient).

Optogenetic perturbation of either PV^+^ or SOM^+^ cells during sound presentation impaired performance of the task (Fig. 6B,J), reducing the sensitivity of the mice to the stimulus (Fig. 6C,K, PV::ArchT *p* = 0.0046, SOM::ArchT *p* = 0.0051, Wilcoxon signed-rank test). Moreover, this reduction in sensitivity did not differ between the manipulation of PV^+^ cells and the manipulation of SOM^+^ cells (*p* = 0.321, Wilcoxon rank-sum test). The observed reduction in performance could not be fully explained by visual distraction from the laser, as the majority of mice had larger deficits in sensitivity when the laser was directed at the auditory cortex rather than directed away (77% of PV::ArchT mice, 80% of SOM::ArchT mice), and there was no significant correlation between the changes in sensitivity for each of these laser location conditions (Fig. 6F,N; PV::ArchT *p* = 0.609, *r* = 0.157; SOM::ArchT *p* = 0.546, *r* = 0.218; linear correlation coefficient).

We found that the reduction in sensitivity from perturbation of inhibitory interneurons was, for both cell types, explained by a reduction in hit rate (Fig. 6D,L, PV::ArchT *p* = 0.033, SOM::ArchT *p* = 0.0051, Wilcoxon signed-rank test), while false alarm rate was not significantly affected (Fig. 6E,M; PV::ArchT *p* = 0.861, SOM::ArchT *p* = 0.878, Wilcoxon signed-rank test). In PV::ArchT animals, neither hit rate nor false alarm rate was correlated between conditions where the laser was directed to the auditory cortex and directed away (Fig. 6G,H; hit rate *p* = 0.657, *r* = −0.136; false alarm rate *p* = 0.981, *r* = −0.0073; linear correlation coefficient). In SOM::ArchT animals, we observed some degree of correlation between the laser directed to the auditory cortex and directed away, reaching statistical significance for hit rate but not false alarm rate (Fig. 6O,P; hit rate *p* = 0.0494, *r* = 0.633; false alarm rate *p* = 0.0725, *r* = 0.590; linear correlation coefficient). Nonetheless, in the majority (70%) of SOM::ArchT animals, hit rate was more strongly affected when the laser was directed at the auditory cortex, suggesting that visual distraction cannot fully account for the effects on hit rate. Our observation that these correlations barely reach statistical significance in one mouse line but not the other could be the result of individual variation in distractibility, or in the amount of spurious light blocked by each implant. Overall, however, these results indicate that, contrary to our original hypothesis regarding different roles for distinct inhibitory neuron classes during signal detection, perturbing PV^+^ or SOM^+^ cell activity results in comparable effects on behavioral output in the signal-in-noise detection task describe here. These effects were apparent in individual behavioral sessions, and plotting the sensitivity index for each session illustrated the reliability of the effect even under the expected variability in behavior from session to session (Fig. 7).

**Figure 7:**
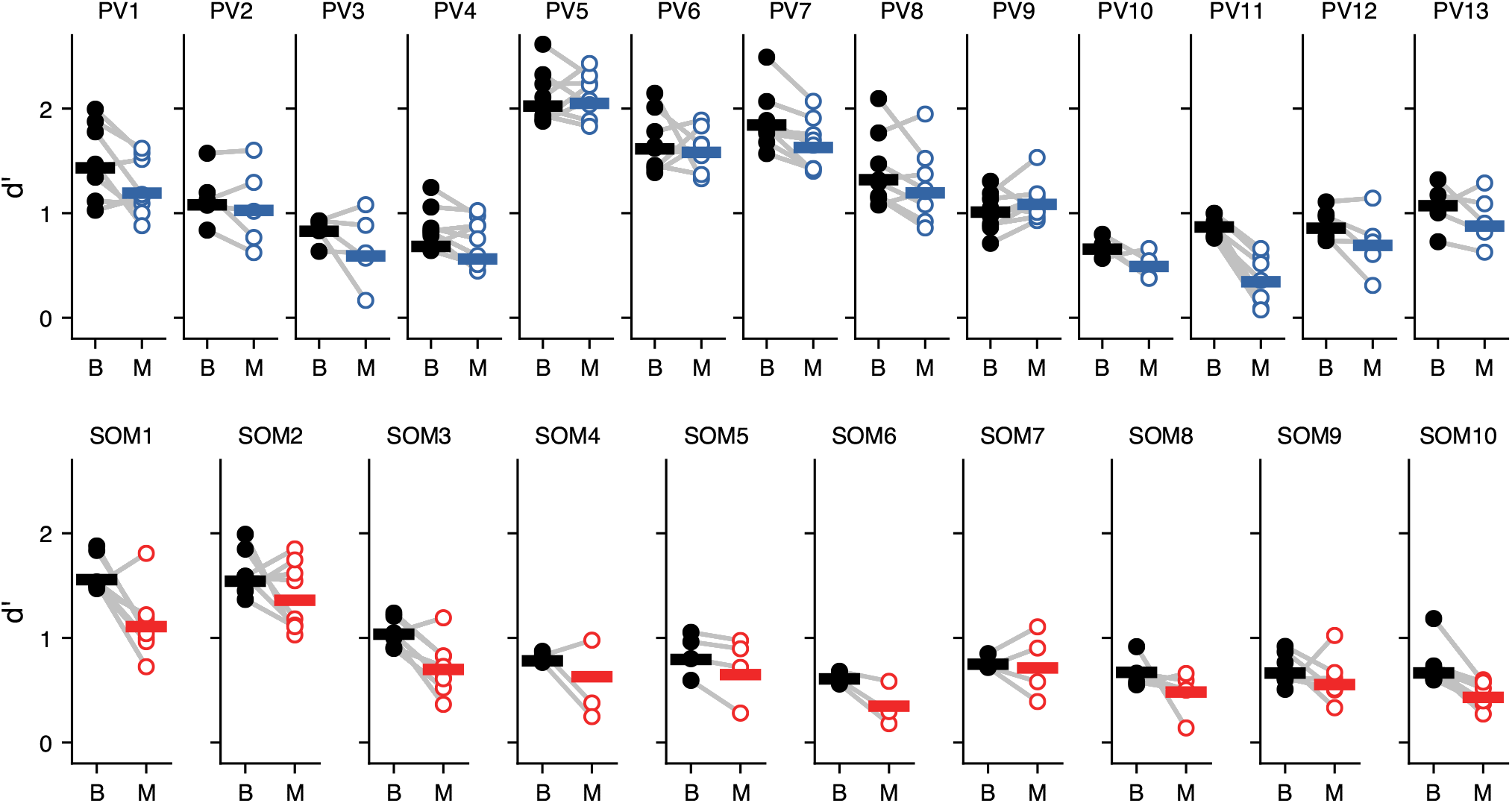
Effects of disrupting auditory cortical inhibition on each behavioral session. Signal detection performance (*d*′) during each behavioral session (dots) compared to the pooled performance across all trials (horizontal lines) for each PV::ArchT mouse (PV1-PV13) and each SOM::ArchT mouse (SOM1-SOM10). B: baseline (laser off); M: manipulation (laser on).

To test whether perturbation of either PV^+^ or SOM^+^ cells from the auditory cortex resulted in any motor deficits during the task, we quantified changes in the timing of the animals’ motor output during laser trials. Perturbation of SOM^+^ cells, but not PV^+^ cells, increased the amount of time mice spend sampling sound, although the effect sizes were very small in both cases (Fig. 8A,E; PV::ArchT *p* = 0.0652, 1% change, SOM::ArchT *p* = 0.005, 3.7% change, median change of 3 ms). The effects on sampling time in PV::ArchT animals were strongly correlated between the conditions where the laser was directed at the auditory cortex and directed away (Fig. 8B, *p* = 0.0017, *r* = 0.780, linear correlation coefficient), suggesting that any effect on sampling time that results from PV^+^ cell perturbation may be a result of distraction by light. In contrast, there was no correlation between these laser location conditions in SOM::ArchT animals (Fig. 8F, *p* = 0.901, *r* = 0.0453, linear correlation coefficient), suggesting that the small effects observed from SOM^+^ cell perturbation are not simply the result of visual distraction by the laser. Furthermore, the time mice took to reach the reward ports was minimally affected by perturbation of PV^+^ cells (Fig. 8C, *p* = 0.0029, 1.8% change) and did not reach statistical significance in SOM::ArchT mice (Fig. 8G, *p* = 0.203, 1.4% change). There was no correlation in time-to-reward between the conditions where the laser was directed at the auditory cortex *vs*. away in either mouse strain (Fig. 8D,H; PV::ArchT *p* = 0.901, *r* = 0.0383; SOM::ArchT *p* = 0.470, *r* = 0.259; linear correlation coefficient). These results indicate that perturbation of PV^+^ or SOM^+^ cell activity does not result in substantial motor deficits, suggesting that the effects on behavioral output are likely perceptual in nature.

**Figure 8:**
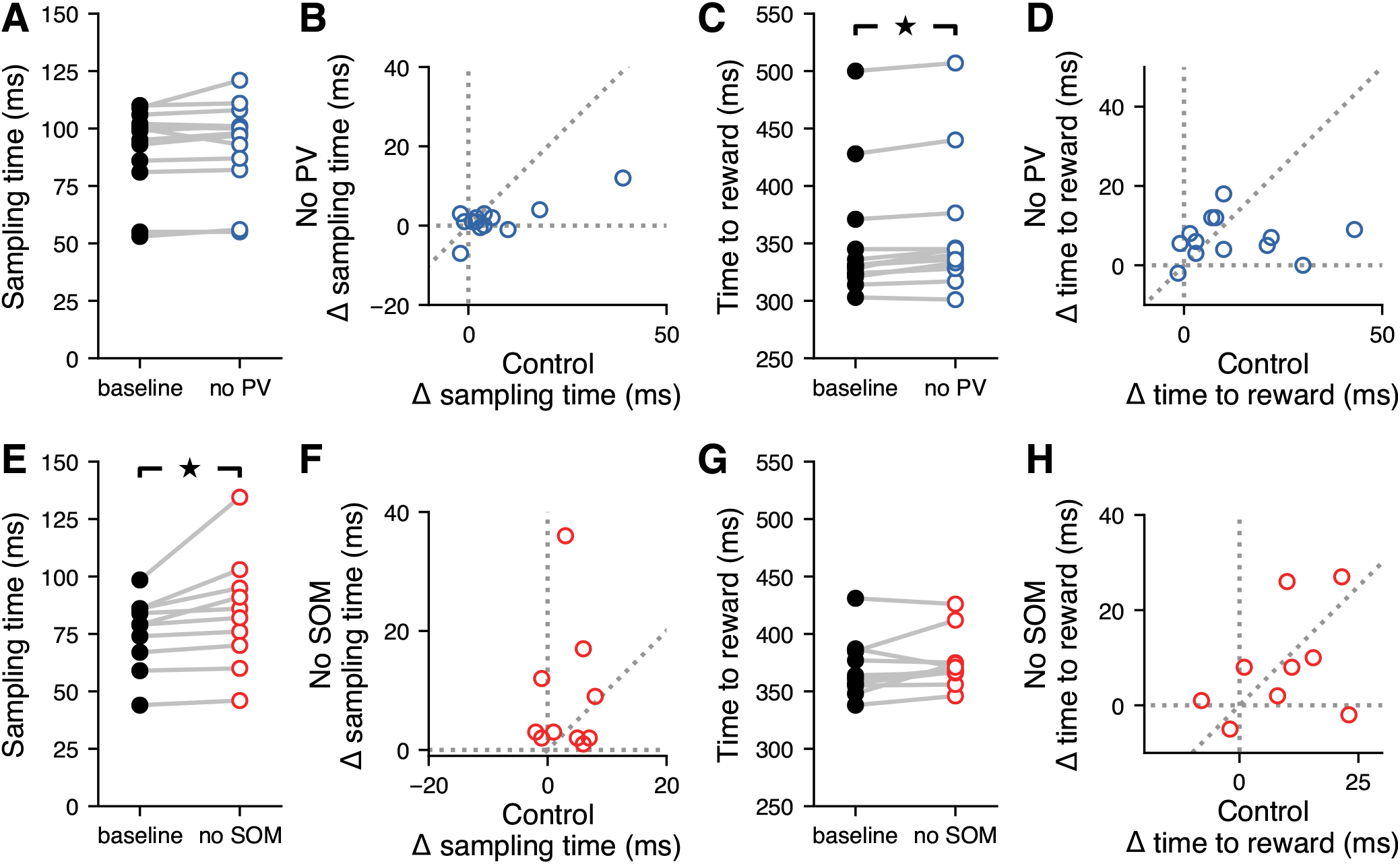
Inactivation of distinct inhibitory neuron types did not affect timing of behavior. **A**, Inactivating auditory cortical PV^+^ cells during the signal detection task did not affect the time spent sampling the sound in the center port (*p* = 0.0652, Wilcoxon signed-rank test). **B**, The effect of inactivating auditory cortical PV^+^ cells (No PV) on the time spent sampling the sound in the center port was correlated with the effect of visual distraction (control) on sampling time (*p* = 0.0017, *r* = 0.780, linear correlation coefficient). **C**, Inactivating auditory cortical PV^+^ cells during the signal detection task led to a small (1.8%) but statistically significant increase in the time spent obtaining a reward (*p* = 0.0029, Wilcoxon signed-rank test). **D**, The effect of inactivating auditory cortical PV^+^ cells on the time spent obtaining a reward was not correlated with the effect of visual distraction on the time spent obtaining a reward (*p* = 0.901, *r* = 0.0383, linear correlation coefficient). **E**, Inactivating auditory cortical SOM^+^ cells during the signal detection task significantly increased the time spent sampling the sound in the center port, leading to a median increase in sampling time of 3.7% (*p* = 0.005, Wilcoxon signed-rank test). **F**, The effect of inactivating auditory cortical SOM^+^ cells (No SOM) on sampling time was not correlated with the effect of visual distraction (control) on sampling time (*p* = 0.901, *r* = 0.0453, linear correlation coefficient). **G**, Inactivating auditory cortical SOM^+^ cells during the signal detection task did not affect the time spent obtaining a reward (*p* = 0.203, Wilcoxon signed-rank test). **H**, The effect of inactivating auditory cortical SOM^+^ cells on the time spent obtaining a reward was not correlated with the effect of visual distraction on the time spent obtaining a reward (*p* = 0.470, *r* = 0.259, linear correlation coefficient).

## Discussion

In this study, we investigated how different sources of inhibition within the auditory cortex contribute to the perception of a pure tone stimulus immersed in background noise. Previous studies have found population-level noise-invariant representations of masked auditory stimuli in the auditory cortex (Christison-Lagay et al., 2017), and discovered that the auditory cortex plays a role in mediating signal detection and discrimination in noisy environments (Sollini and Chadderton, 2016; Christensen et al., 2019). Consistent with these studies, we found that optogenetically silencing the auditory cortex during sound presentation resulted in mice becoming less likely to report the presence of narrowband signals, providing further evidence that the auditory cortex plays an important role in identifying behaviorally relevant signals in noisy acoustic environments. Interestingly, silencing the auditory cortex also reduced the rate at which false alarms occurred, but did not affect the timing or direction of movement. This suggests that the observed effects on behavior were perceptual in nature rather than motor, and that outputs from the auditory cortex are partially responsible for the perception of hallucinatory stimuli. Furthermore, inactivation of either PV^+^ or SOM^+^ cells during sound presentation created a deficit in performance specifically by decreasing hit rates, indicating that perturbing inhibitory auditory cortical circuits disrupts the outputs that carry information about the presence of signals.

The optogenetic techniques used here for reversibly silencing cortical circuits via activation of cortical PV^+^ cells, or for inactivating specific inhibitory neurons, have been validated extensively by others (Sachidhanandam et al., 2013; Glickfeld et al., 2013; Weible et al., 2014a,b; Natan et al., 2015). Our physiological recordings from a PV::ArchT mouse also indicate that perturbing PV-expressing interneurons impacts the responses of neurons across the auditory cortex. However, the interpretation of any experiments that use optogenetics must take into account the possibility that expression patterns may differ slightly across animals, altering the effects on each individual. These caveats may be minimized by performing electrophysiolgical recordings on each animal used for behavior analysis, a challenging task not included in our experiments, and therefore a limitation of our study.

Contrary to the expectation that detecting loud stimuli would rely less on auditory cortical circuits than detecting quiet stimuli, we observed comparable effects for signals with low and high SNR when inactivating the auditory cortex. Therefore, the observed limited ability of mice to separate signals from noise during perturbation of cortical circuits does not seem to be explained by a partially inactivated auditory cortex. Instead, under the assumption that detecting loud sounds does not require as much of the auditory cortex to be intact, our results are better explained by the proposition that we are fully silencing the auditory cortex, but some residual processing is preserved in other auditory areas.

Our results indicate that activating and inactivating PV^+^ cells have very similar effects on behavior. While a decrease in sensitivity (*d*′) is an expected result of any perturbation of normal auditory cortical activity, perturbing PV^+^ cells in either direction resulted in the same effect on hit rate, raising the question of why opposite manipulations of PV^+^ cells would result in similar effects on signal detection. Two mechanisms contribute to the ability to detect signals in noise: suppression of responses to broadband noise and amplification of responses to narrowband signals (Atiani et al., 2009). The former has been shown to be mediated in part by cortical inhibition (Lakunina et al., 2020), while the latter may be performed by the auditory cortex or inherited from upstream areas. The auditory cortex could use a combination of these mechanisms to determine the presence or absence of signals and pass this information to downstream areas. Thus, silencing the auditory cortex entirely (through the activation of PV^+^ cells) would lead to a decrease both in hit rate and false alarm rate because the neural signals triggering the perception of auditory stimuli are failing to travel to downstream areas. In contrast, inactivation of PV^+^ cells may disrupt the auditory cortical circuits needed to suppress responses to broadband noise, leading to a decrease in hit rate (as with activation of PV^+^ cells), but not affect the circuits amplifying narrowband signals, leading to no change in false alarm rate. Consistent with these ideas, we observed that activation of PV^+^ cells led to a decrease in false alarm rate, while inactivation of PV^+^ cells did not have an effect on false alarm rate.

Our experiment relied on inactivating PV^+^ cells with ArchT to determine the effect this cell type plays on separating auditory signals from noise. Under some circumstances, optogenetic inactivation of PV^+^ with ArchT may be ineffective at reducing PV^+^ firing rates due to recurrent network effects causing a paradoxical increase in both PV^+^ and excitatory cell responses (Moore et al., 2018). Still, an increase in excitatory activity, which we observed in electrophysiolgical recordings in PV::ArchT animals, is indicative of a perturbation of the normal function of PV^+^ cells in cortical circuits, and the behavioral effects presented here can be interpreted as the result of disrupting the normal function of PV^+^ cells during detection of auditory signals. An additional concern when manipulating neural activity with optogenetics is that activation and inactivation of a particular cell type may not produce entirely opposite effects on physiological responses due to the nonlinearities produced by thresholds or saturation of neural activity (Phillips and Hasenstaub, 2016). While these caveats may partly explain our observation that activation and inactivation of PV^+^ cells result in similar decreases in hit rate, opposite effects on neural responses may not necessarily result in opposite effects on behavior. As described in the previous paragraph, it is plausible that both silencing the auditory cortex entirely (through the activation of PV^+^ cells) and disrupting inhibitory auditory cortical circuits (through the inactivation of PV^+^ cells) could result in similar effects on hit rates.

This study sought to determine whether PV^+^ and SOM^+^ cells have different contributions to the ability of mice to detect a behaviorally relevant signal immersed in broadband noise. Previous studies have shown that PV^+^ and SOM^+^ cells have different contributions to lateral inhibition and surround suppression (Adesnik et al., 2012; Kato et al., 2017; Lakunina et al., 2020), processes that we hypothesized would be important to the suppression of responses to broadband noise. Despite this, we found that inactivation of either PV^+^ or SOM^+^ cells had, on average, similar effects on behavioral output, in both cases leading to a reduction in hit rate and no effect on false alarm rate. This may indicate that, if background subtraction occurs in the auditory cortex, the sources of inhibition underlying this mechanism are not split along the broadly-defined gene-expression categories we studied. It is also possible that PV^+^ and SOM^+^ cells play different roles in the implementation of the mechanisms that give rise to noise-invariant responses, but the perturbation of their activity has widespread effects on the auditory cortex which degrade performance in auditory signal detection tasks. Spontaneous activity in the auditory cortex is modulated to increase the signal-to-noise ratio during behavioral engagement in signal detection tasks (Buran et al., 2014; Carcea et al., 2017), and disruption of the peripheral auditory system hinders signal detection by increasing signal-to-noise ratio in the auditory cortex (Resnik and Polley, 2021). It is therefore plausible that perturbation of either PV^+^ or SOM^+^ cells increases baseline activity in the auditory cortex (Natan et al., 2015) and worsens the signal-to-noise ratio of responses to auditory stimuli, consistent with our behavioral results.

Our results reveal that inhibitory auditory cortical circuits are important for making decisions about the presence or absence of behaviorally relevant acoustic signals in a noisy background. The relation between the distinct effects PV^+^ and SOM^+^ cells have on the responses of excitatory cells and the potentially different roles these inhibitory cells play in the perception of sounds in noise remains an open question.

## Acknowledgements

This research was supported by the National Institute on Deafness and Other Communication Dis- orders (R01DC015531), and the Office of the Vice President for Research & Innovation at the University of Oregon.

